# Spherical harmonic based noise rejection and neuronal sampling with multi-axis OPMs

**DOI:** 10.1101/2021.12.22.473837

**Authors:** Tim M Tierney, Stephanie Mellor, George C O’Neill, Ryan C Timms, Gareth R Barnes

**Affiliations:** Wellcome Centre for Human Neuroimaging, UCL Queen Square Institute of Neurology, University College London, London WC1N 3AR, U.K.

**Author notes:** **Corresponding author**: Tim Tierney, Wellcome Centre for Human Neuroimaging, UCL Queen Square Institute of Neurology, University College London, 12 Queen Square, London WC1N 3AR, UK; E.

## Abstract

In this study we explore the interference rejection and spatial sampling properties of multi-axis Optically Pumped Magnetometer (OPM) data. We use both vector spherical harmonics and eigenspectra to quantify how well an array can separate neuronal signal from environmental interference while adequately sampling the entire cortex. We found that triaxial OPMs have superb noise rejection properties allowing for very high orders of interference (L=6) to be accounted for while minimally affecting the neural space (2dB attenuation for a 60-sensor triaxial system). To adequately model the signals arising from the cortex, we show that at least 11^th^ order (143 spatial degrees of freedom) irregular solid harmonics or 95 eigenvectors of the lead field are needed to model the neural space for OPM data (regardless of number of axes measured). This can be adequately sampled with 75-100 equidistant triaxial sensors (225-300 channels) or 200 equidistant radial channels. In other words, ordering the same number of channels in triaxial (rather than purely radial) configuration gives significant advantages not only in terms of external noise rejection but also minimizes cost, weight and cross-talk.

## 1 Introduction

Optically Pumped Magnetometers (OPMs) and Superconducting Quantum Interference Sevices (SQUIDS) both measure the brain’s neuromagnetic field (Baillet, 2017; Xia et al., 2006). However, the measured signal differs in both its amplitude and spatial information content. These differences arise because OPMs can be placed directly on the scalp and therefore sample higher spatial frequencies of the brain’s magnetic field at greater magnitude (Iivanainen et al., 2021). In theory these higher spatial frequencies should result in better spatial resolution, but if their information is to be represented without any form of signal aliasing more dense arrays or custom sensor layouts are required (Ahonen et al., 1993; Beltrachini et al., 2021; Tierney et al., 2020; Vrba & Robinson, 2002).

One powerful approach to quantifying the degree of higher spatial frequency content in MEG has been to use the Signal Space Separation (SSS) method (Taulu & Kajola, 2005). This approach has the added benefit that it can simultaneously model an array’s ability to reject environmental interference. This is because SSS provides a model for both the neural data and environmental interference. The models for the neuronal data and the interference respectively are the gradients of irregular and regular spherical harmonics. The irregular harmonics tend to zero at coordinates far from coordinate system’s origin (outside the sensor array) and are thus useful for modelling magnetic fields coming from inside the sensor array (brain signals). The regular harmonics become zero at the origin (inside the array) and are therefore useful for modeling magnetic fields that come from outside the array (interference).

An open question for OPM recordings is how many orders of spherical harmonics are required to model the environmental interference and the neuronal data. With regards to interference, our previous work has suggested that the limited sensor numbers in typical OPM arrays can result in the attenuation of neuronal signal if too high a spatial order of interference is selected (Tierney et al., 2021). Furthermore, it is worth considering whether the ability of OPMs to measure in more than one direction affects the selection of the spherical harmonic model order for the neuronal signal and magnetic interference. This is an important consideration as previous work on cryogenic systems has led to diverging approaches to system design. For example, work on (SQUID based) cryogenic systems has suggested that a two-layer, sensor array (comprising 400 sensors) measuring in more than 1 direction should achieve shielding factors greater than 60dB using SSS (Nurminen et al., 2010). It has been demonstrated empirically even adding a small number of tangential channels improves shielding factors (Nurminen et al., 2013). However, other authors have argued that optimal system design (for maximizing SNR) consists of 1^st^ order radial gradiometers combined with 3^rd^ order synthetic gradiometers (Fife et al., 1999; Vrba & Robinson, 2002).

Optimal system design for OPM recordings is less clear as one has the added issues of optimizing for wearability and subject movement (Boto et al., 2018). For example, optically pumped gradiometers offer promising noise cancellation properties (Limes et al., 2020; Sheng et al., 2017), yet as larger sensor baselines are used, one may compromise wearability. The tradeoff is therefore to shorten the baseline (Nardelli et al., 2020) but this will result in lower depth sensitivity (Hämäläinen et al., 1993; Vrba & Robinson, 2002). Software approaches for magnetometers not reliant on (fixed) reference arrays such as beamformers (van Veen & Buckley, 1988), SSS (Taulu & Kajola, 2005) or Signal Space Projection (Uusitalo & Ilmoniemi, 1997) are thus quite attractive. Recent work on OPMs has suggested that beamformers have their interference control improved by multi-axis recordings (Brookes et al., 2021) allowing for movement in excess of 1m to be made (Seymour et al., 2021). With these issues in mind, it is worth exploring to what extent a single layer of vector OPMs can separate brain signal from magnetic interference and how this interacts with sampling density.

There are many diverging viewpoints and methods to optimize neuronal sampling with OPMs. One could use different definitions of optimality or information content (Beltrachini et al., 2021; Iivanainen et al., 2021), including minimization of aliasing (Tierney et al., 2020) or seeking to minimize correlation between sources (Boto et al., 2016). One extensive exploration of radial oriented OPMs suggest that between 177 and 276 sensors are required for sufficient sampling (Iivanainen et al., 2021). The broad estimate of requisite number of sensors arises due to the use of different basis sets (eigenbasis and spatial frequency basis) for the estimation of the spatial degrees of freedom in the data. The SSS basis set can also provide an estimate of the spatial degrees of freedom in the data and we compare this estimate with the estimate from the eigenbasis.

Throughout this work we rely on the SSS framework to explore the issues raised. We chose to use SSS as a basis as it provides a unifying theoretical framework for both neuronal sampling and environmental interference rejection. All of its interference rejection properties can be derived theoretically once the geometry of the MEG array is known. It also allows one to provide an upper bound on the number of spatial degrees of freedom in OPM data. We can therefore theoretically explore the interactions of multi-axis recordings and varying sampling densities while modeling neural data and magnetic interference. We expect these results to be useful for those wishing to design OPM arrays for MEG experiments.

## 2 Theory

### 2.1 SSS as a model of brain signal and interference

SSS represents the MEG signal (*H*) as a linear combination of the gradients of spherical harmonics (*Y*_*lm*_(*θ, φ*)) with coefficients *α*_*lm*_ and *β*_*lm*_. The full formulation for the magnetic field in spherical coordinates (*r, θ, φ*) is

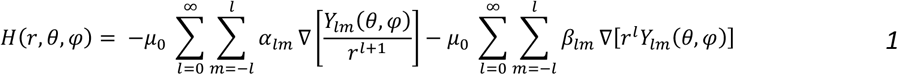

where *μ*_0_ is the magnetic permeability. The first term represents the neural space (with the gradient of irregular spherical harmonics) while the second term represents the interference space (with the gradient of regular spherical harmonics). When modelling MEG recordings, each set of basis functions is truncated to a maximum value of *l*. This value is chosen in order to represent sufficient variance in the signal or sufficiently model environmental interference. The number of columns in each basis set is 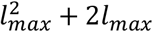, where *l*_*max*_ is the order of the harmonic used. The spherical harmonic basis functions (*Y*_*lm*_(*θ, φ*)) are complex functions. For a more intuitive representation of the harmonics, we replace them with real valued harmonics, as in previous work (Mellor et al., 2021). In cartesian coordinates (*x, y, z*) these harmonics 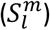 are known as the solid harmonics and are defined as follows:

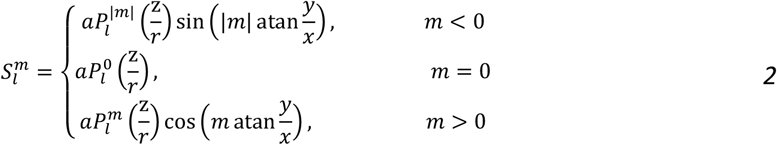

where the associated Legendre polynomial 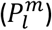 has the following form:

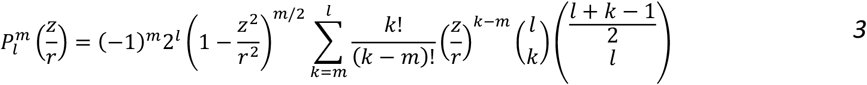

with

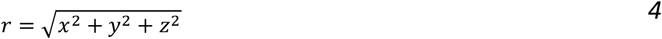

and

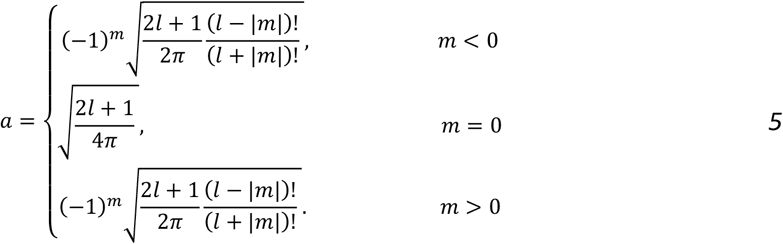

For ease of notation we refer to the basis set representing the neural space and interference space as *A* and *B* respectively.

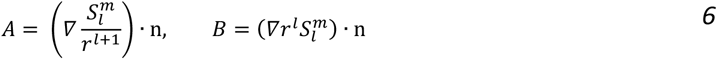

With n being the unit vector representing the senor’s sensitive axis and · represents the dot product. We provide an explicit form for *A* and *B* in Appendix I and code to create these harmonics is made publicly available at https://github.com/tierneytim/OPM/blob/master/spm_opm_vslm.m

### 2.2 The orthogonality of interference and neuronal data

For the model of the interference (*B*) to be practically useful, it needs to share minimal variance with the lead fields (*L*). This is not guaranteed for every array design. To asses this we measure the lead field variance attenuation when the interference term is regressed from the data. This regression step can be summarized in one projection as follows

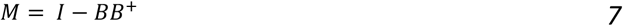

where *B*^+^ is the pseudoinverse. We can now summarize the variance attenuation for each brain area in decibels (dB)

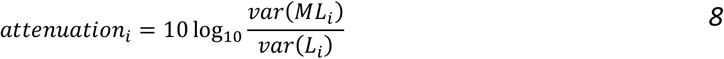

where *var*(*L*_*i*_) is defined as

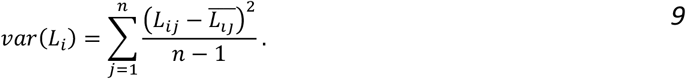

The indices *i, j* refer to the magnetic field produced by the *i*^*th*^ brain area (in this case a vertex on a cortical mesh) at the *j*^*th*^ sensor for *n* sensors. 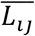 refers to the arithmetic mean for the *i*^*th*^ brain area across sensors. Now we have a metric for every brain area that summarizes how much variance is lost for a given regular solid harmonic order of *B*. We can also calculate these metrics for any sensor array or number of measurement axes. This is crucial for establishing robustness of a given array design to environmental interference.

### 2.3 The order of harmonics required to model the neural space

As OPMs sample higher spatial frequencies one would assume they would require higher orders of harmonics in matrix *A* to fully model the neural data. We arbitrarily describe the data as fully modelled when 99% of signal variance is explained in greater than 95% of brain regions. Similarly, to section 2.2 we can measure the variance explained (*VE*_*i*_) as

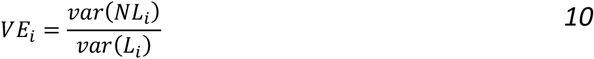

where

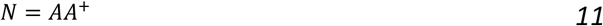

The order of harmonics required to model the neuronal data is important as using sufficiently high order harmonics would allow us to bandlimit (in terms of the spatial frequencies represented by the harmonics) the OPM data. Any interference or noise outside this bandlimit would then be suppressed. It also allows us to come up with an upper bound for the number of independent samples in the data.

### 2.4 Assessing the efficiency of the neural model

The more efficiently a given model can represent the neural data the more interference can be rejected. An “efficient” model would have fewer parameters than there are measurements. As a benchmark to compare the use of solid harmonics to model the neural data, we also consider eigenvectors (*V*) of the lead field covariance (*C*) matrix to model the neural data.

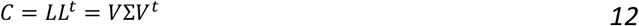

Where Σ represents the covariance matrix’s eigenvalues. Comparing the basis set *V* and *A* in terms of their efficiency is of interest as algorithms such as DSSP (Cai et al., 2019; Sekihara et al., 2016) project the data onto the lead field eigenvectors. The software for MEG/EEG analysis, SPM, also performs this step as a preprocessing procedure for source reconstruction (Friston et al., 2008; Lopez et al., 2014). As *V* is an orthogonal matrix the projector (*O*) that projects the data onto this basis is simply

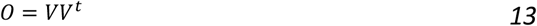

We can then once again calculate the variance explained as follows

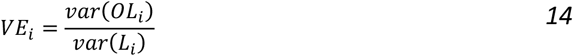

In summary, we have now derived two subspace definitions which can be used to model neuronal signal, one based on solid harmonic gradients (*A*, eq. 6) and one based exclusively on the MEG system lead-fields (matrix *V* eq. 12). If we use the exact same threshold for when we consider the brain fully modelled (>99% signal power in >95% of brain regions) we can compare the relative efficiency of both models (*V*and *A*) in representing neural data. We can then explore how both these models change as a function of sampling density and number of measured axes.

### 2.5 Considerations for non-spherical sampling

We also consider the implications of the using spherical harmonic models of neural data in non-spherical sampling situations. As one places sensors closer to the brain the relative influence of different sensors may change based on their distance to the origin (of the sensor coordinate system). This is because as the harmonic orders get higher in the matrix *A*, the dependency on the distance to the origin increases. If every sensor has the same distance to the origin (spherical sampling) this does not matter, but if sensors are not equidistant to the origin some sensors may have excessive influence on the modelling. This process can be captured formally with the statistical concept of Leverage.

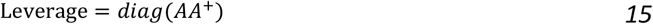

Each Leverage value tells us how influential a given observation is on the model. More formally it is the rate of change of the model with respect to the data. We can make this measure relative by dividing it by its mean value. Now each leverage value tells us how influential a data point is relative to the average data point. It is computed as follows:

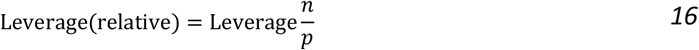

where *n* is the number of sensors and *p* is the number of parameters of the model.

## 3. Methods

### 3.1 Lead field generation

OPMs were modelled as point magnetometers displaced 6.5mm from the scalp. The brain mesh used to generate these lead-fields was the MNI canonical mesh available in SPM12 with 8196 vertices. The separation between vertices is approximately 5 mm on average. The orientation of the source was defined by the surface normal of the cortical mesh at that location. The forward model was the Nolte single shell model (Nolte, 2003). The sensors were placed on the scalp surface with separations of 85 to 15mm in steps of 5mm. The sensor placement algorithm is described elsewhere (Tierney et al., 2020). For each level of sensor spacing we simulated single axis, dual axis and triaxial sensors, generating 3 lead-field matrices per sensor spacing.

### 3.2 The Orthogonality of interference and multi-axis OPM recordings

We generated the first three orders of the regular solid harmonics in cartesian coordinates. The lead field attenuation (as documented in section 2.2) was then calculated for each order, at each level of spatial sampling for single, dual and triaxial recordings respectively. We also calculated the lead field attenuation for orders L=1 to 12 for single, dual and triaxial systems comprising of 60 and 400 sensors. This second analysis is intended to compare the limits of interference rejection in a realistic wearable array to an ideal, but impractical, array.

### 3.3 Order of harmonics required to model the neural space

We generated the first 15 orders of the real irregular solid harmonics in cartesian coordinates for the densest sampling (15mm separation). The variance this basis set explained in the lead fields was calculated for single, dual and triaxial systems, as described in section 2.3. For comparison we also calculated the variance explained for magnetometers displaced 24mm from the scalp (to represent a cryogenic, SQUID based system). At this point we also estimate the impact of non-spherical sampling on the model of the neural data. We computed the leverage as described in section 2.5 and compared the sensors’ influence on the neuronal model to its distance to the origin (of the sensor coordinate system) at different orders of spherical harmonics (L=1, L=6, L=12). For comparison we also calculated these same metrics on a sphere. For simplicity of presentation we only examine one sensitive axis.

### 3.4 Assessing the efficiency of the neural model

The efficiency of the model to represent the neural data (the number of basis vectors required) is compared against the eigenvectors of the lead field which also form a compact basis set for describing the neural space (Iivanainen et al., 2021). To assess how many eigenvectors are required to model the neural space we compute the variance explained as in section 2.4. We do this for all steps of spatial sampling for, single, dual and triaxial measurements.

### 3.5 Software

Software required to generate the vector spherical harmonics described in this paper is made freely available on the first author’s GitHub page (https://github.com/tierneytim/OPM). The key function is spm_opm_vslm. Examples and tests can also be found on GitHub (https://github.com/tierneytim/OPM/blob/master/testScripts/testVSM.m).

## 4. Results

### 4.1 The Orthogonality of Interference and multi-axis OPM recordings

Figure 1 (A, B, C) shows the expected lead field attenuation when regressing regular solid harmonics of increasing order (L=1, L=2, L=3) from OPM data for single axis, dual axis and triaxial sensors. As expected, as the number of sensors is increased there is less risk of attenuating sensitivity to neuronal sources. Interestingly, all sensor types (triaxial, dual axis and single axis) converge rapidly to an expected signal loss at all investigated orders. For greater than 30 sensors (90 channels) the expected signal loss is lower than 1dB for triaxial sensors even at high harmonic order (L=3). This is in contrast to single axis sensors which see greater than 15dB attenuation at this point. For L=3, This same information is represented spatially on the brain (Figure 1D) for radial, dual and triaxial sensors. Ultimately, triaxial sensors allow for suppression of more spatially complex interference patterns with minimal risk of brain signal loss.

**Figure 1.**
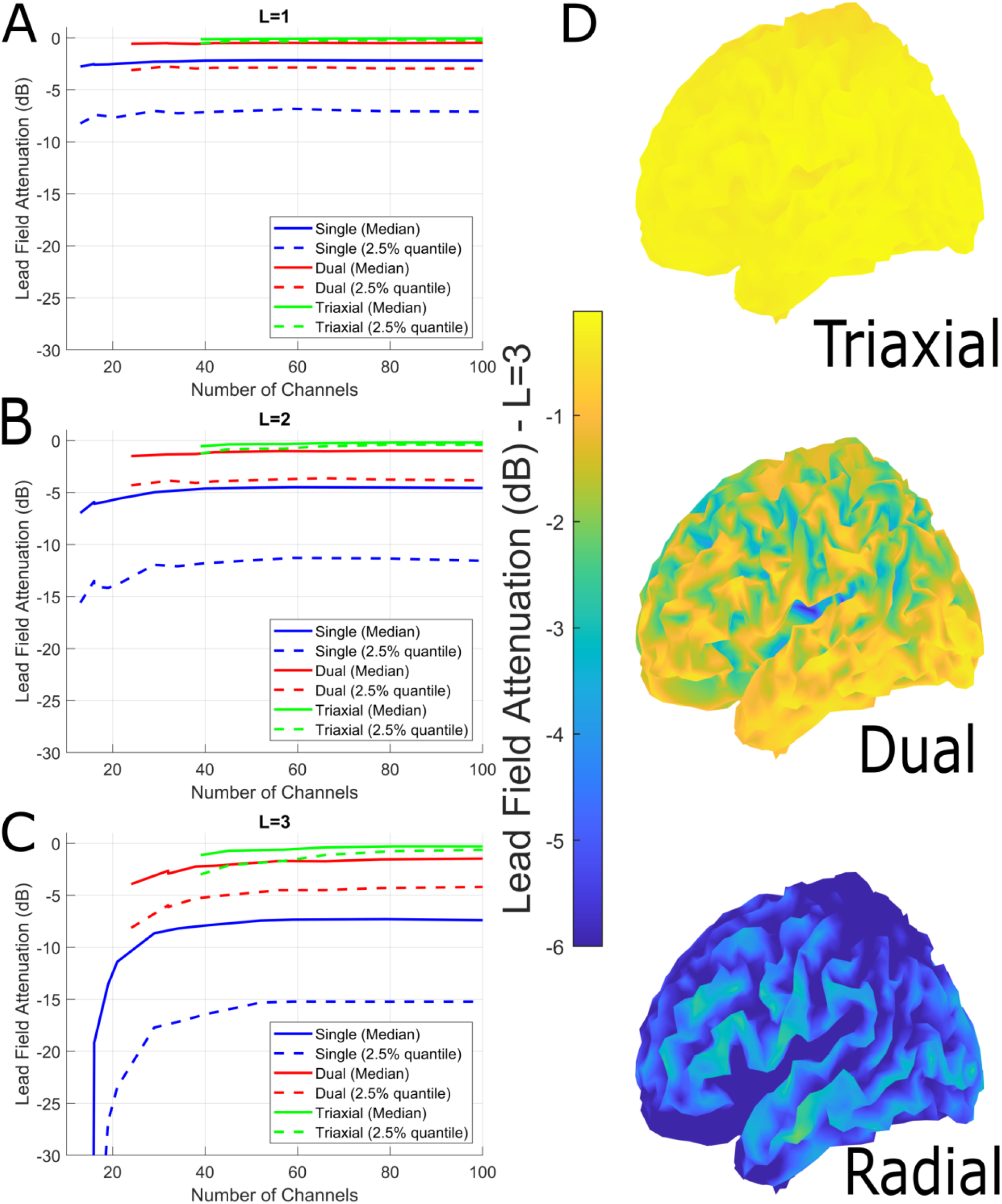
The orthogonality of interference and multi-axis OPM recordings. A, B and C show the expected lead field attenuation in decibels (y axis) when regressing regular solid harmonics of increasing order (L=1, L=2, L=3) from OPM data for single axis (blue), dual axis (red) and triaxial (green) sensors. Solid lines show the median expected signal loss and dashed lines show the signal loss from the worst-affected top 2.5% of brain regions (worst case scenario). The x-axis indicates the number of channels. D shows the spatial distribution of the expected signal loss for L=3, for radial, dual-axis and triaxial sensors. To summarize the figure, multi-axis recordings allow the removal of more complex environmental interference whilst preserving the neuronal signal. Importantly this is partly a factor of channel number but heavily determined by geometry as lower numbers of triaxial channels have less attenuation then large numbers of single axis or dual axis sensors (eg compare 60 tri-axials to 60 dual-or 60 single-axis sensors).

In Figure 2A and 2B we explore the statistical and physical limit of using high order models of interference on OPM data. Unsurprisingly, as the number of regressors (*order*^2^ + 2 × *order*) approaches the number of channels the attenuation grows rapidly for all systems representing the statistical limit of interference control (Figure 2A). In Figure 2B when the number of sensors is large relative to the number of regressors the lead field attenuation is determined by the spatial similarity of the magnetic fields generated by the brain and the interference space (representing a physical limit of using higher order models). However, one can see that for even a 60-sensor system the order of harmonics that one could remove from the data without exceeding 3dB of (neuronal space) attenuation for a triaxial system is greater than 6.

**Figure 2.**
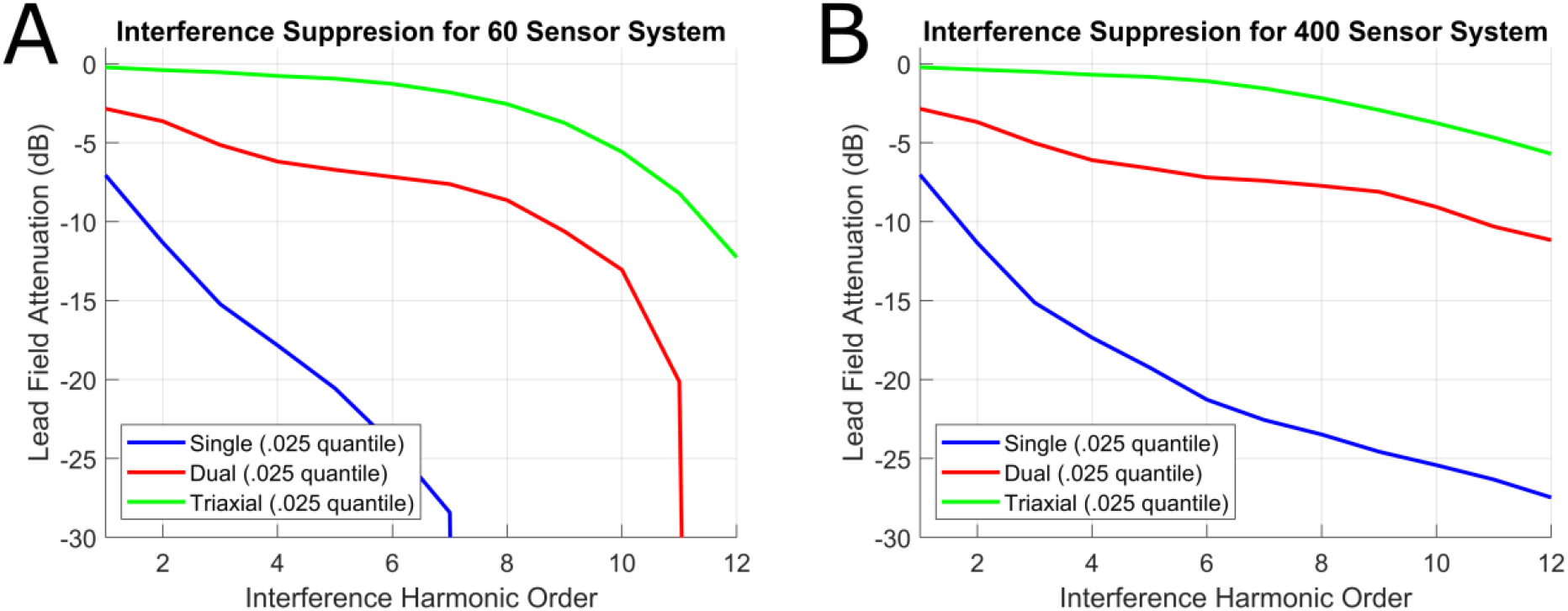
Statistical and physical limits of interference suppression for a 60 (A) and 400 channel (B) OPM system. **A**. The 2.5% quantile lead field attenuation (decibels) for a 60-sensor system measuring single, dual and triaxial channels respectively. In other words, for a triaxial system, one could remove ∼10 orders of external interference at the price of attenuating neuronal signals from 2.5% of the cortex by 5dBs. Triaxial systems offer clear advantages over dual and single axis systems in terms of minimal attenuation at high harmonic order. As the number of regressors (order^2^ + 2 × order) approaches the number of channels the attenuation grows rapidly for all systems (representing the statistical limit of interference control). The same results are plotted in B for a 400-sensor system. In this case, the number of regressors never approaches the number of channels and we do not see the sudden attenuation of lead field variance at higher orders. In this case the lead field attenuation is representative of the physical limit of using higher order models.

### 4.2 Order of harmonics required to model the neural space

Figure 3 shows the order of irregular solid harmonics required to explain OPM data (6.5mm scalp offset point magnetometers) and SQUID data (24mm scalp offset point magnetometers) for single, dual and triaxial systems. These simulations were performed on the data with the highest degree of spatial sampling (15mm separation). Encouragingly, for the SQUID data (which only differs from the OPM data in scalp offset), the saturation of the harmonics (achieving 99% variance explained for 95% of brain regions) occurs at the same harmonic order (8) as in previous research (Taulu & Kajola, 2005). The OPM data saturates at L=11 implying there are at most 143 spatial degrees of freedom (see discussion for comparison with existing literature). Perhaps surprisingly, the results of Figure 3 also imply that multi-axis measurements provide no new spatial information concerning the brain’s activity when compared to single axis measurements. However, this is due to the large number of sensors used in the simulation and will be explored further in the next section.

**Figure 3.**
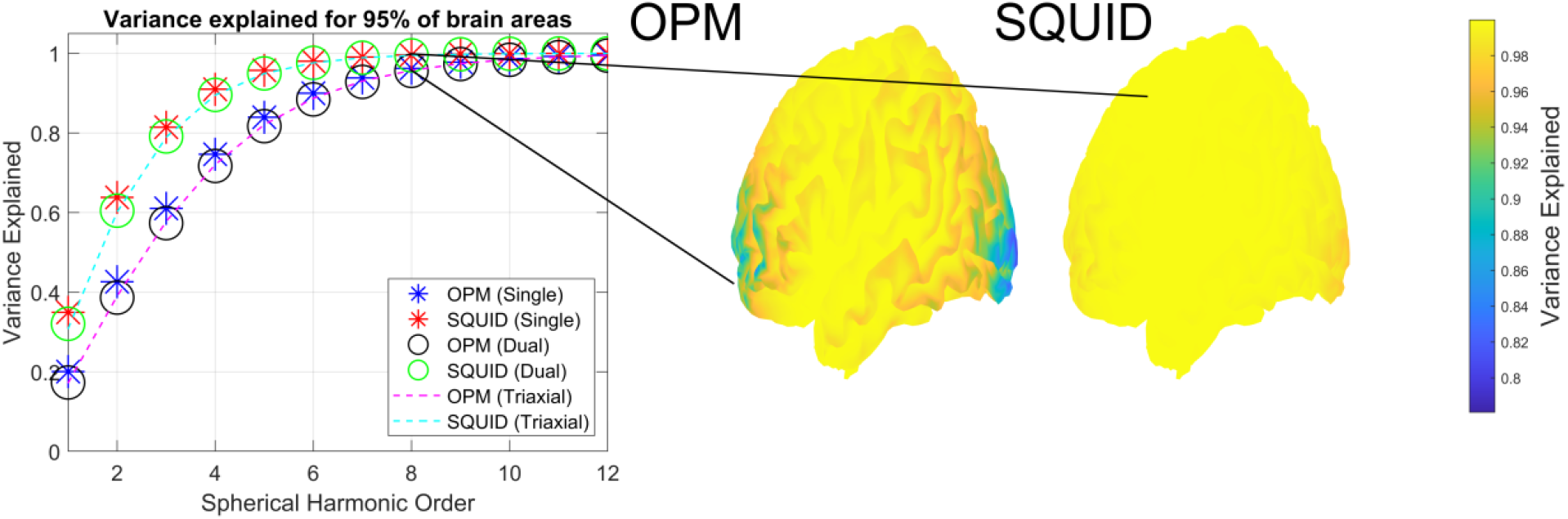
Order of irregular harmonics required to model OPM data. The y axis shows the variance explained for a given harmonic order (x axis) for OPMs and SQUIDS for single, dual and triaxial sensors (stars, circles and dashed lines repsectively). On the right we show the spatial profile of the variance explained across the brain for both OPMs and SQUIDS at L=8.

If we examine where the irregular solid harmonics poorly explain the brain data we note a spatial profile. As an example, at L=8 (Figure 3 right) it would appear the harmonic model only explains 80% of the variance in areas at the front and back of the head in OPM data. This effect is largely explained by the sensors that are most distant from the origin of the coordinate system having the least influence on the model (Figure 4). This phenomenon is more pronounced for data with higher spatial frequency content (such as OPM data) as higher harmonic orders (L=6, L=12) have many more highly influential sensors than at lower orders (L=1). Interestingly, in the case of spherical sampling, each sensor is equally influential on the model regardless of spatial frequency content.

**Figure 4.**
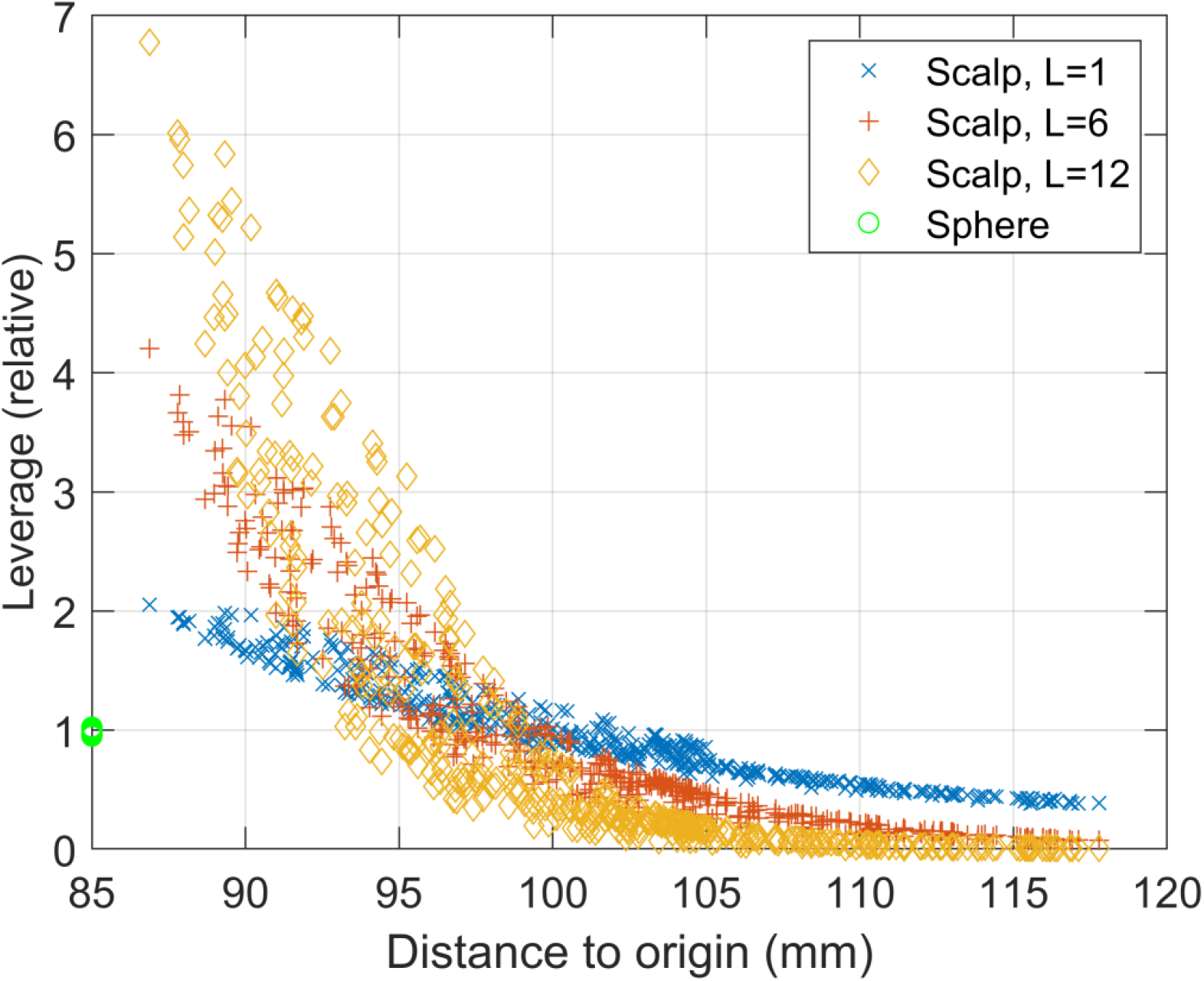
Sensor influence on irregular harmonic model of brain activity. Sensors displaced far from the origin of the coordinate system (x axis) have less relative leverage (y axis) than sensors closer to the coordinate system origin. This effect is stronger at higher orders of harmonics. Essentially, sensors displaced far from the origin have minimal impact on the model at high orders. This analysis is performed for both scalp-based sampling and sampling on a sphere. Note that although sampling on a sphere provides the most equitable impact of sensors on the model it is a special case where regular and irregular harmonics become highly correlated (perfectly correlated in the case of single axis radial sensors) rendering the separation of brain signal from interference highly challenging.

### 4.3 Information content of multi-axis sensors at lower sampling densities

The results of the previous sections would suggest that while multi axis recordings provide a much better model of magnetic interference they provide no new spatial information about the neural signal when compared to a large arary of single axis sensors. We explore this phenomenon by examining the eigenspetra of the leadfields. We argue that sufficient sampling of the neural space has occurred when additional sensors do not increase the number of eigenvectors that contribute substantial variance to the lead fields. This is seen in Figure 5, where the number of eigenvectors required to explain 99% of variance in 95% of brain areas saturates at lower sensor numbrs for triaxial sensors when compared to dual or single axis sensors. In fact, the figure shows that there are diminishing returns (in terms of characterising neuronal signal) for triaxial systems after 75 sensors (∼225 channels). This is not the case for an equally distributed radial system which does not saturate until 150-200 sensors (or channels) while the triaxial system completely saturates at 100 sensors (300 channels). Figure 5B shows these same data in terms of channel rather than sensor count and we see that radial only systems can be constructed with lower channel counts to sample the same (neuronal) information.

**Figure 5.**
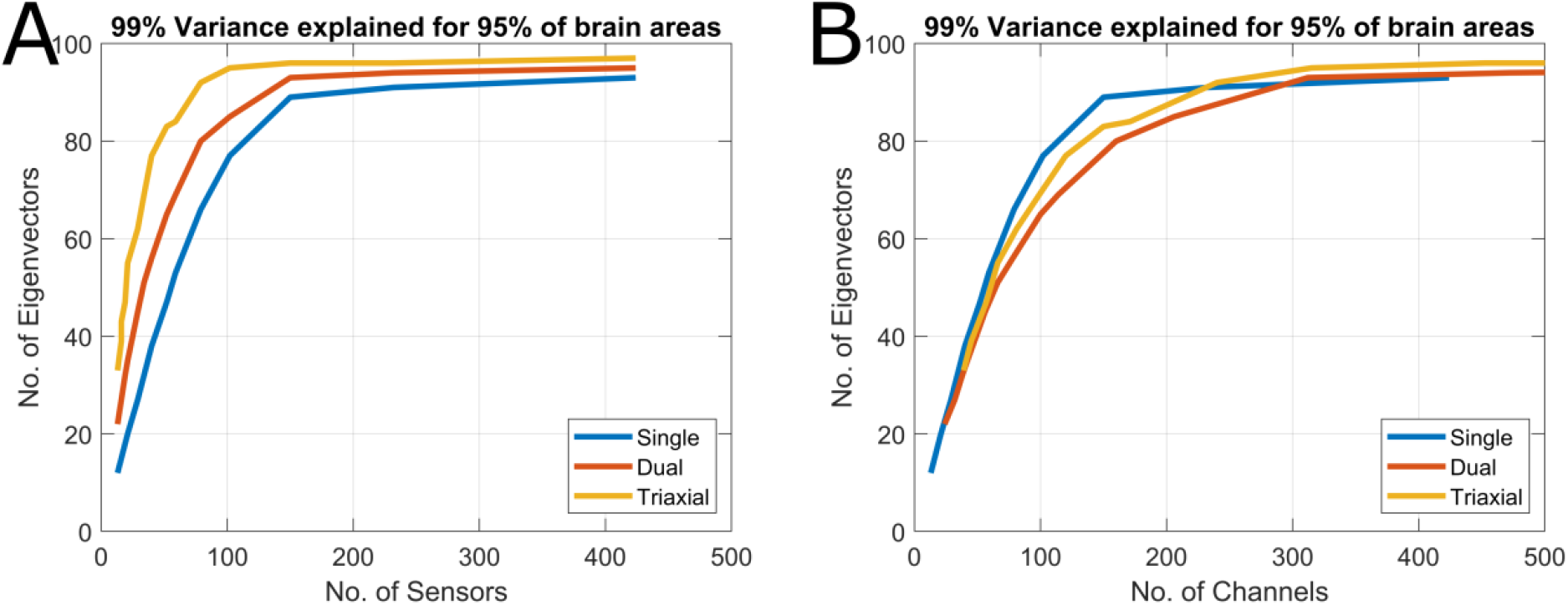
Eigenspoectra of single, dual and triaxial OPM system. A. The figure graphs the number of sensors (x-axis) vs the number of eigenvectors (y-axis) to ahchive 99% variance explained in >95% of brain region. When the number of eigenvectors stops changing as a funcion of sensor number, one has fully sampled the neural space. B shows the same information but the x axis is changed to channel count. The comparison of these two figures celarly shows that while triaxial systems allow for sampling the neuronal signal with lower sensor counts (A) radial only systems can be constructed with lower channel counts to sample the same information (B).

## 5. Discussion

By using vector spherical harmonics and eigenpectra as a theoretical basis we have explored the interference rejection and spatial sampling properties of single, dual and triaxial OPM data. We found that triaxial OPMs have superb noise rejection properties allowing for very high orders of interference (L=6) to be accounted for while minimally affecting the neural space (2dB attenuation for a 60-sensor triaxial system). The neural space was efficiently modelled by both irregular solid harmonics (L=11, number of triaxial/dual/radial channels =143) and by the eigenvectors of the lead fields (number of triaxial sensors and eigenvectors < 100). We now discuss the implications for system design of these findings.

Spherical harmonics are commonly used to model the brain and magnetic interference with the SSS method (Clarke et al., 2020; Nurminen et al., 2010; Taulu et al., 2005; Taulu & Simola, 2006). Our work here reveals that for OPM data the default parameters derived for SQUID data should change for OPM data. In SQUID data the neural space is modelled with harmonics of order 8 and the interference is modelled with harmonic of order 3. For OPM data the neural space should be modelled with at least harmonics of order 11 regardless of the number of axes measured. As an interesting side note we found that if one were to truncate the model before L=11 (Figure 3), the front and the back of the brain are poorly modelled. This is because as one places sensors closer to the brain, a sensor’s ability to influence the model becomes strongly dependent on its distance to the origin of the coordinate system (Figure 4).

When modelling interference, the appropriate order strongly depends on the number of measurement axes. In short, radial only designs will perform poorly even at low harmonic order but triaxial systems will perform well at very high orders. Importantly, this effect is not driven by differing number of channels between single, dual and triaxial systems. We can see clearly in Figure 1 that regardless of how many radial channels are utilized, the lead field attenuation will always be greater than that due to 20 triaxial sensors (60 channels). In triaxial systems, the limit for how high an order can be used will be determined by the number of regressors in the interference model approaching the number of data points (Figure 2). While it is clear that having more axes improve noise rejection (Brookes et al., 2021; Nurminen et al., 2010), rejection performance is also improved by the sensors being closer to the brain. Proximity to the brain shifts the neural spatial frequency content higher (Figure 3) and magnetic interference (which is spatially low frequency at a distance) becomes progressively more orthogonal to the neural signal. As such, on-scalp systems should (theoretically) have better noise rejection properties than off-scalp systems.

With regards to sampling, we have built upon previous work which has shown that custom arrays can be designed to model a particular brain region (Beltrachini et al., 2021; Iivanainen et al., 2021; Tierney et al., 2020). In the current work we show that 100 equidistant triaxial sensors are sufficient to model the whole brain. In fact, there are diminishing returns in having more than 75 triaxial sensors (Figure 5A). In terms of channel number, the triaxial arrangement is not as efficient as a purely radial array (Figure 5B). Essentially, while the vector components of a triaxial system do add independent information, the information gain per channel is smaller than it is for a radial only design. However, in addition to the enhanced noise rejection properties there are practical considerations. As the manufacturing of triaxial OPMs negligibly affects their weight and cost, it may be beneficial to design a sparser triaxial array than a dense single axis array to optimize subject comfort and array wearability. Furthermore, the average sensor spacings are increased from 22mm (for single axis only) to 32mm for triaxial. Increasing the distance between sensors by 50% will have the added benefit of reducing the effects of sensor cross-talk (Nardelli et al., 2019) by a factor > 2.

With regards to limitations we are assuming that a triaxial sensor can be constructed without compromising performance. It has been shown that tangential components may be more affected by volume currents (Iivanainen et al., 2017) and we do not investigate the implications here or of the impact of different forward models (Stenroos et al., 2014; Stenroos & Sarvas, 2012). It is also worth considering that measuring multiple axes also often occurs a small increase in white noise (Osborne et al., 2018). However, even if triaxial sensors come with additional white noise burden, we note that environmental interference such as vibration induced noise can exceed 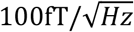 easily (Tierney et al., 2021). As such, we are (currently) often limited by interference and not white noise. To mitigate this increased white noise, one could project the triaxial system on to either the eigenvectors of the lead fields (matrix *V* in section 2) or onto the irregular solid harmonics (matrix *A* in section 2).

We expect this white noise mitigation to be greater for projecting onto the eigenvectors of the lead field than on to irregular solid harmonics (assuming white noise is spread equally among the regressors of *V* and *A*) as *V* required < 100 regressors to represent the neural signal whereas *A* required at least 143 regressors. The compact way in which *V* represents the neuronal signal is important as projecting the data onto *V* is a preprocessing step applied in all SPM source reconstruction algorithms (Friston et al., 2008; Lopez et al., 2014) and as a first stage in preprocessing for the noise rejection algorithm DSSP (Cai et al., 2019; Sekihara et al., 2016).

There also are many other algorithms used for interference suppression that are not considered here. We chose to use SSS as a basis because all of its interference rejection properties can be derived theoretically once the geometry of the MEG array is known. It also allows one to provide an upper bound on the number of spatial degrees of freedom in OPM data. Other data driven techniques such as ICA (Vigârio et al., 2000), tSSS (Taulu & Hari, 2009) or the canonical correlation step in DSSP (Cai et al., 2019) are more difficult to consider because they are data driven. However, a full consideration of the applicability of these techniques for suppressing interference in OPMs is warranted. As a final point we have also not considered whether the positions and orientations of dual axis sensors could be further optimized to have performance similar to triaxial systems. In the current work we have first simulated a radial sensor and then arbitrarily chosen the second axis. In principle the first axis need not be radial and the second could be optimized.

With these limitations in mind, we have shown theoretically that triaxial OPM sensors are capable of separating signal from inference (with higher spatial frequency content) with minimal risk of attenuating brain signal when using regular solid harmonics. Furthermore, sparser arrays can be constructed with triaxial sensors than radial sensors (at the cost of increased channel count). These findings all suggest that future systems based around triaxial arrays could allow for minimization of cost, weight and interference while maximizing the system’s sensitivity to neural data in sparse arrays.

## Appendix I: Real, Cartesian, Vector Spherical Harmonics

Here we provide explicit expressions for the matrices *A* and *B* introduced in section 2.

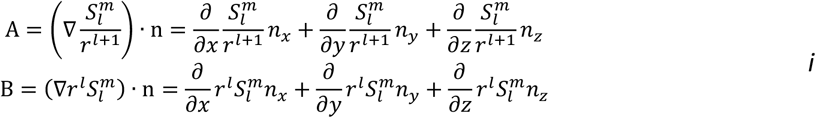

where *n*_*x*_, *n*_*y*_, *n*_*z*_ refer to the components of the unit vector (*n*) representing the sensors sensitive axis. We introduce the variable *t* to refer to any of the axes, *x, y, z* and calculate the partial derivatives as below.

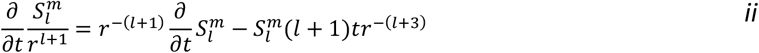

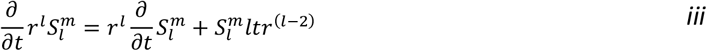

The only unknown in the above equation is 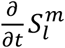 which can be expressed as follows in Cartesian coordinates

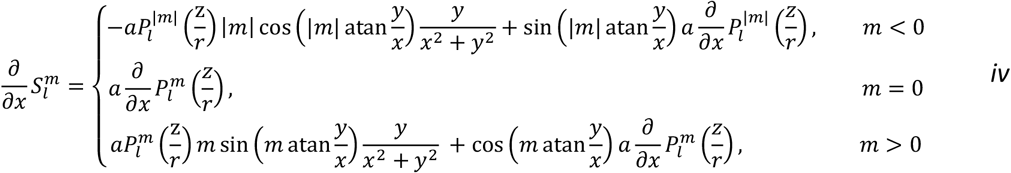

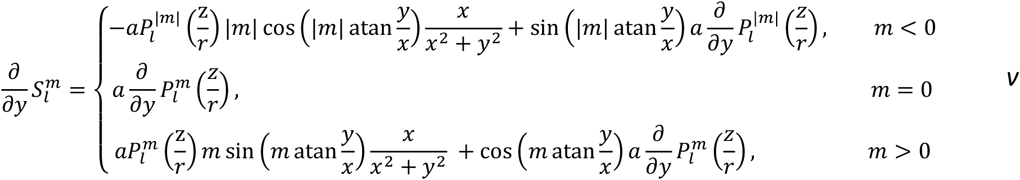

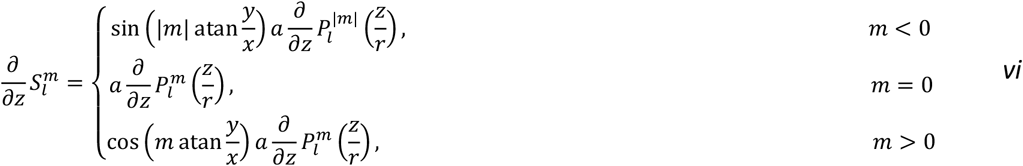

Where

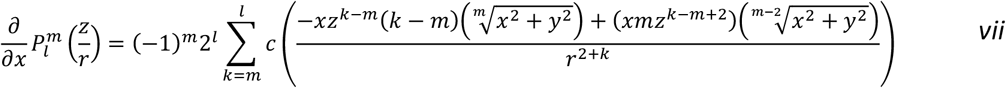

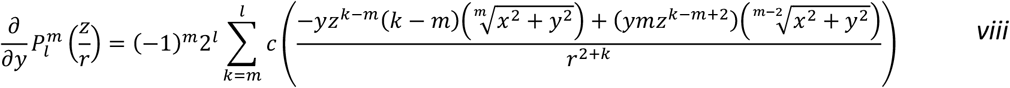

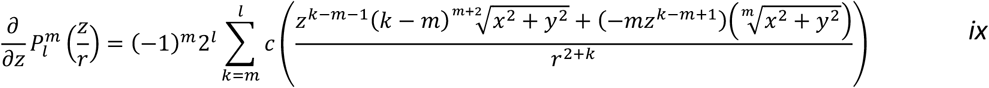

With *c* defined as follows

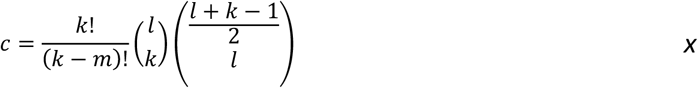

## Acknowledgements

This work was supported by a Wellcome collaborative award to GRB, Matthew Brookes and Richard Bowtell (203257/Z/16/Z, 203257/B/16/Z). SM was funded through the EPSRC-funded UCL Centre for Doctoral Training in Medical Imaging (EP/L016478/1) and GO through EPSRC (EP/T001046/1) funding from the Quantum Technology hub in sensing and timing (sub-award QTPRF02). The Wellcome Centre for Human Neuroimaging is supported by core funding from the Wellcome Trust (203147/Z/16/Z). We also would like to thank Vishal Shah and his team at QuSpin for their support throughout the development of our OPM system.

